# Systemic orchestration of cell size throughout the body: Influence of sex and rapamycin exposure in *Drosophila melanogaster*

**DOI:** 10.1101/2023.01.11.521715

**Authors:** Ewa Szlachcic, Anna Maria Labecka, Valeriya Privalova, Anna Sikorska, Marcin Czarnoleski

## Abstract

Along with differences in life histories, metazoans have evolved vast differences in cellularity, involving changes in the molecular pathways controlling the cell cycle. The extent to which the signalling network systemically determines cellular composition throughout the body and whether tissue cellularity is organized locally to match tissue-specific functions are unclear. We cultured genetic lines of *Drosophila melanogaster* on food with and without rapamycin to manipulate the activity of TOR/insulin pathways and evaluate cell-size changes in five types of adult cells: wing and leg epidermal cells, ommatidial cells, indirect flight muscle cells, and Malpighian tubule epithelial cells. Rapamycin blocks TOR multiprotein complex 1, reducing cell growth, but this effect has been studied in single cell types. As adults, rapamycin-treated flies had smaller bodies and consistently smaller cells in all tissues. Regardless, females eclosed with larger bodies and larger cells in all tissues than males. Thus, differences in TOR activity and sex were associated with the orchestration of cell size throughout the body, leading to differences in body size. We postulate that the activity of TOR/insulin pathways and their effects on cellularity should be considered when investigating the origin of ecological and evolutionary patterns in life histories.

## Introduction

Within modern vertebrates, the fish *Schindleria brevipinguis* matures with a body mass approximately 284,000,000-fold lighter than that of the blue whale *Balaenoptera musculus* [1,2]. Apparently, along with differences in body plans and life histories, organisms have evolved enormous differences in body size, which differentiate phylogenetic branches, populations [3–6] and sexes [7,8]. Moreover, changes in adult size are also commonly involved in the plasticity of organisms’ developmental responses to environmental conditions [9–12]. Such high variability in the size of organisms has long inspired scientific debate [13–16], mainly because the size achieved at maturity is a fundamental determinant of Darwinian fitness [17,18], with far-reaching consequences for physiological and ecological processes, e.g., the energy budget, mortality, competition, and niche breadth [13,17]. Mechanistically, this variability involves changes in cell numbers and size as well as the amount of extracellular components, but we usually do not know the role of each of these mechanisms or their fitness consequences [8,13,18]. Emerging evidence suggests that cell size does not remain constant, changing both with the developmental environment [12,19–23] and over evolution, differentiating populations and species [19,24]. There is also direct evidence that changes in cell size are a component of plasticity- and evolutionary-related changes in adult size [7,19,21,25–28]. Undoubtedly, the number and size of the cells that make up an organism have consequences for organismal performance [29–33], but it is unclear whether the cellular structure of tissues is systemically organized throughout the metazoan body or whether it is arranged locally to suit tissue-specific functions. A clear answer to this fundamental question is unavailable largely because previous studies have rarely focused on cell-size variance among organisms and even then have tended to target single cell types [10,21,34–37]. The systemic orchestration of cell size throughout the body may be an evolutionarily conserved characteristic [38], which is partially supported by studies of invertebrates [8,12,20,39,40], vertebrates [24,41] and plants [42].

Research is only just beginning to understand how cells sense and regulate their size, with studies focusing on cell-cycle checkpoints and molecular pathways involved in cell-autonomous and systemic control of cell size [33,43]. On a macroevolutionary time scale, the evolution of cell-cycle control appears to have involved the gradual incorporation of new signalling pathways into the conserved backbone formed by AMP-activated protein kinase (AMPK) and target of rapamycin (TOR), two protein kinases that act as signalling hubs to integrate and exchange information with the entire network of regulatory pathways [44,45]. The emergence of eukaryotes probably occurred after the origin of the TOR pathway, which subsequently became regulated by the insulin pathway with the emergence of animals [44,45]. Thus, there seems sufficient evidence to consider the TOR/insulin pathways as good candidates for the primary regulatory mechanism ensuring systemic orchestration of cell size throughout the animal body. To explore this possibility, we experimentally manipulated TOR activity in fruit flies (*Drosophila melanogaster*) by rearing larvae with or without rapamycin and measuring their body size as well as the cell sizes in five organs of the eclosed adult flies. At the molecular level, rapamycin targets TOR, downregulating the activity of TOR multiprotein complex 1 [46]. Rapamycin is a bacterial antibiotic used as an immunosuppressive drug [47], with recognized anti-ageing potential [48]. Rapamycin administration to *D. melanogaster* larvae has been shown to lead to smaller adult flies and smaller cells [29,49,50], but it remains unclear how, if at all, modulation of TOR activity orchestrates cell sizes in different tissues and organs in the body. Additionally, such systemic control may not occur, as emerging (but still fragmentary) evidence in snails [51], geckos [25] and woodlice [39] indicates irregularities in cell-size changes in different tissues. Importantly, previous studies on the effects of rapamycin in flies have typically focused on larvae directly exposed to rapamycin and examined only single cell types and only cells that lose viability after fly metamorphosis (i.e., larval epidermal cells, which are responsible for the production of exoskeleton elements in adults). Clearly, we are far from possessing a satisfactory understanding of the systemic control of cell size and the involvement of the TOR signalling pathway in this phenomenon.

## Methods

We used 14 genetic isolines from a wild population of *D. melanogaster* established in 2017 (49°58’00.8” N, 20°29’54.1” E) and maintained at the Institute of Environmental Sciences (Jagiellonian University, Kraków, Poland) [29]. Flies were maintained in vials (polyurethane vials with foam plugs) with cornmeal yeast medium (Bloomington Drosophila Stock Center, Bloomington, USA) in 40-ml vials (with 10 ml of food) in the stock population and in 68-ml vials (with 20 ml of food) for the study population. Flies were kept in thermal cabinets (POL-EKO, Wodzisław Śląski, Poland) maintained at 20.5° C, with 70% relative humidity at a 12 h:12 h L:D photoperiod. Transfers prevented generational overlap.

Following our previous approach [29], we produced two consecutive generations under conditions of controlled larval density. Upon each transfer, we placed 10 females and 5 males from each isoline in a vial for 48 h for oviposition. On the second transfer, representatives of each isoline were assigned to two treatments, with oviposition performed in vials with rapamycin-supplemented food or standard food. For rapamycin treatment, we dissolved rapamycin (Alfa Aesar by Thermo Fisher Scientific, Kandel, Germany) in 96% ethanol (Linegal Chemicals, Warszawa, Poland) and added it to standard food at a 1 μM concentration. For the control, the same amount of ethanol was added to the standard food. Flies from the second generation, 1-16 days after eclosion, were anaesthetized with CO2. For each fly (112 per treatment), we measured the distance from the neck edge to the tip of the scutellum (thorax length, mm) under a stereomicroscope (Olympus SZX12, Olympus, Tokyo, Japan). The fly was dissected using a microtome knife and forceps. We took a left wing, left middle leg, and head from two males and two females per isoline (56 flies per treatment); in males, we also dissected the Malpighian tubules. From another two males and two females per isoline (56 flies per treatment), we obtained thoraxes. The legs (ethanol), wings (freezing), heads (methanol) and thoraxes (reagents) were preserved as indicated for further steps; Malpighian tubules were imaged immediately without fixing. We measured (Fig. 1) the size of epidermal cells in the wing (μm^2^) and leg (μm), ommatidial cells (μm^2^), Malpighian tubule epithelial cells (μm^2^), and dorsal longitudinal indirect flight muscle cells (μm^2^). We used a light microscope (Eclipse 80i, Nikon, Tokyo, Japan), camera (Axio Cam MRc5, Zeiss, Oberkochen, Germany) and ZEN software (ver. 2011, ZEISS, Oberkochen, Germany) to image the wings, legs, ommatidia, flight muscles (with bright-field microscopy) and Malpighian tubules (with phase contrast objectives). Supplementary materials provide greater detail on the techniques used.

**Figure 1.**
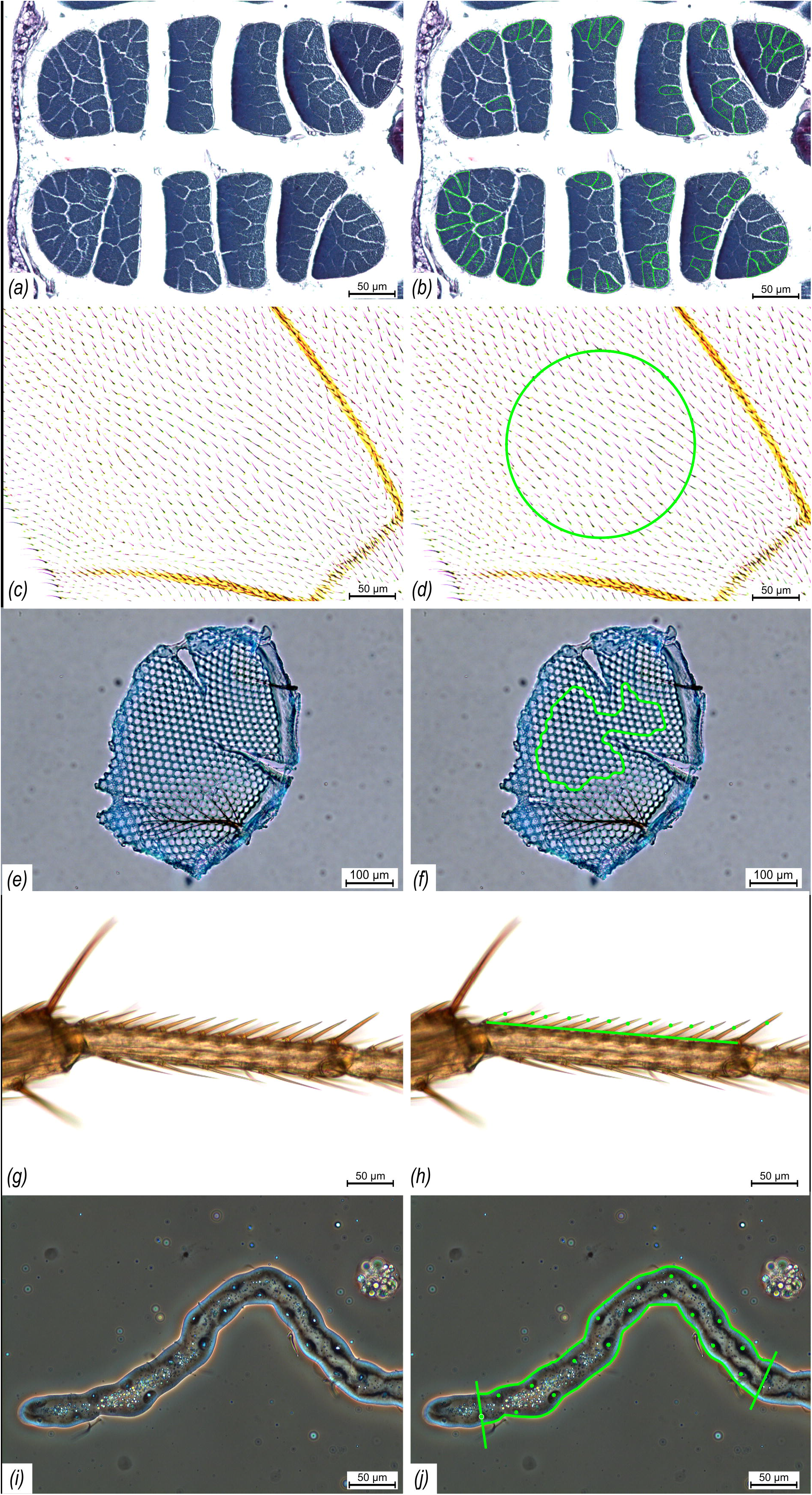
Cell-size measurements in *Drosophila melanogaster* adults. a & b) Thorax: indirect flight muscles (mean cross-sectional area of fibres); c & d) wings: epidermal cells (from trichome number per area unit); e & f) ommatidia: ommatidial cells (mean area of ommatidia); g & h) legs: epidermal cells (from trichome number per length unit); i & j) Malpighian tubules: epithelial cells (from nuclei/nucleoli number per area unit). Left panels show raw images, right panels show images with measurements.

Thorax lengths and cell sizes were analysed with general linear mixed models (GLMMs) in R (v4.0.3) software [52] with the lme4 [53], lmerTest [54] and car [55] packages. Figures were generated with g the gplot2 [56] and emmeans [57] packages as well as Inkscape [58]. To normalize the data distribution, thorax lengths were cube transformed. The models included treatment (rapamycin vs. control) and sex (Malphigian tubules were obtained for males only) as fixed factors and isoline as a random factor.

## Results

GLMMs demonstrated that rapamycin supplementation produced flies with smaller thoraxes (by 6.9% in males and by 4.8% in females; Fig. 2a) and reduced cell sizes in all tissues (Table 1) including in flight muscles (by 12.0% in males and 10.5% in females; Fig. 2b), epidermal cells in wings (by 5.6% in males and 4.7% in females; Fig. 2c), ommatidial cells (by 5.3% in males and 5.0% in females; Fig. 2d), epidermal cells in legs (by 1.6% in males and 1.5% in females; Fig. 2e) and Malpighian tubule epithelial cells (by 36.7% in males; Fig. 2f). For cells in the legs, the results were not significant at p=0.05 but showed a pattern consistent with other tissues. Compared to males, females had larger thoraxes (by 11.8% in control flies and 14.3% in rapamycin flies; Fig. 2a) and larger cells in all cell types (Table 1) including flight muscles (by 14.7% in control flies and 16.7% in rapamycin flies; Fig. 2b), epidermal cells in wings (by 18.0% in control flies and 19.0% in rapamycin flies; Fig. 2c), ommatidial cells (by 6.3% in control flies and 6.6% in rapamycin flies; Fig. 2d) and epidermal cells in legs (by 5.0% in control flies and 5.1% in rapamycin flies; Fig. 2e).

**Figure 2.**
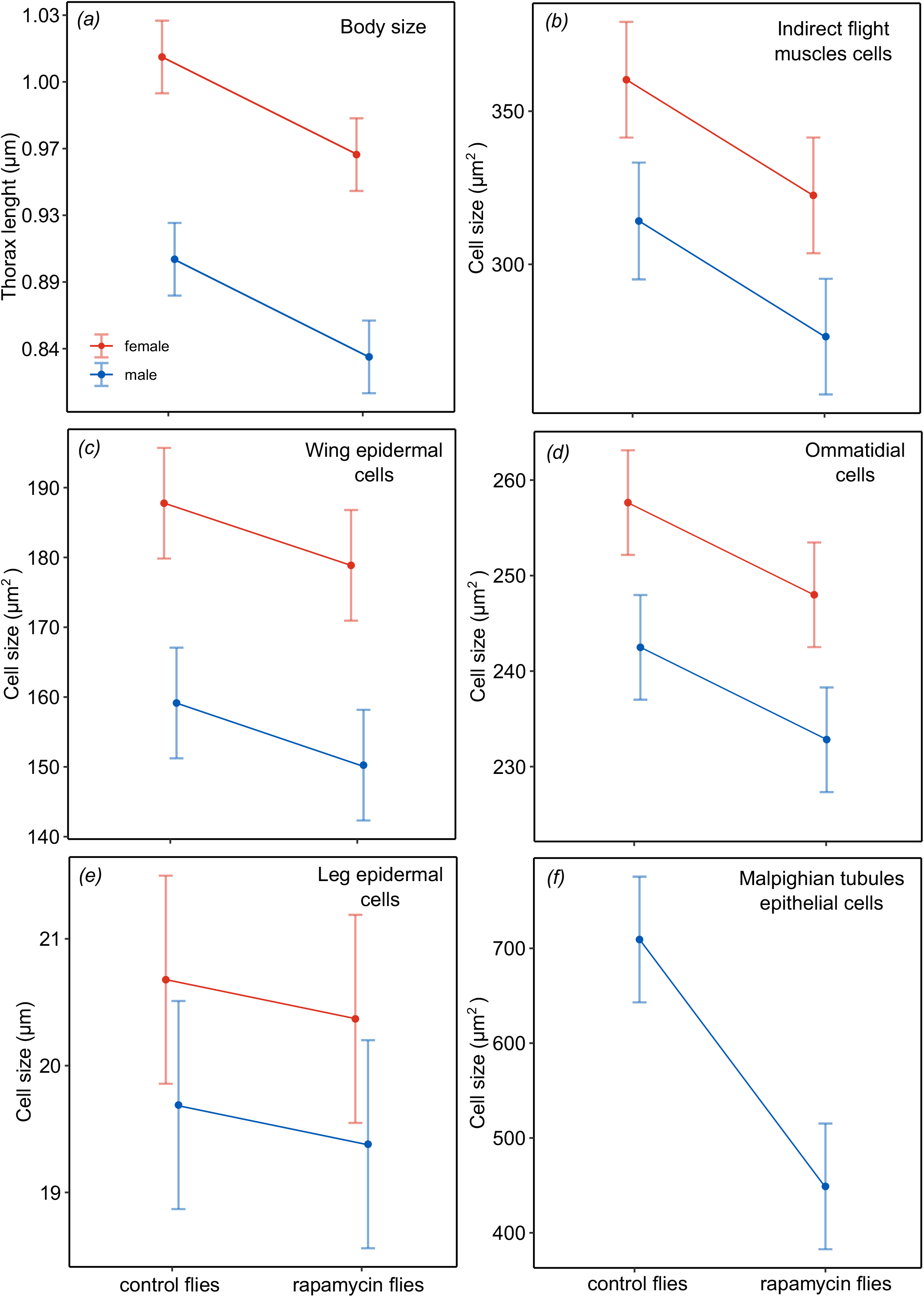
After exposure to rapamycin during development, eclosed *Drosophila melanogaster* had smaller thoraxes (a) and cell sizes in five tissues (b-f; Fig. 1). Modelled means with 95% confidence intervals (Table 1). Thorax length was back-transformed for display.

**Table 1.** Comparison of adult *Drosophila melanogaster* after feeding diets with and without rapamycin.

| Effects in GLMMs | Body size: thorax length |  | Cell size: thorax muscle |  | Cell size: wings |  | Cell size: ommatidia |  | Cell size: legs |  | Cell size: Malpighian tubules |  |
| --- | --- | --- | --- | --- | --- | --- | --- | --- | --- | --- | --- | --- |
|  | N=224 |  | N=111 |  | N=112 |  | N=109 |  | N=112 |  | N=56 |  |
|  | t | P | t | P | t | P | t | P | t | P | t | P |
| Intercept (control female) | 41.1 | <0.0001 | 38.3 | <0.0001 | 50.0 | <0.0001 | 97.7 | <0.0001 | 52.7 | <0.0001 | 21.7 | <0.0001 |
| Treatment (rapamycin) | -10.1 | <0.0001 | -3.9 | 0.0002 | -4.7 | <0.0001 | -7.3 | <0.0001 | -1.3 | 0.1890 | -5.9 | <0.0001 |
| Sex (male) | -21.0 | <0.0001 | -4.8 | <0.0001 | -15.1 | <0.0001 | -8.6 | <0.0001 | -4.2 | <0.0001 |  |  |

## Discussion

Rapamycin supplementation of *D. melanogaster* larvae reduced cell sizes in all five organs assessed in adult flies, with females consistently exhibiting larger cells than males; this finding indicates strong systemic orchestration of cell sizes throughout the body. Specifically, depending on the organ and sex of an individual, rapamycin induced a 1.5–36.7% reduction in cell size; females had cells 5–19% larger than those of males. The systemic cell-size changes were involved in the origin of body-size differences, such that smaller adult flies (rapamycin/male individuals) had consistently smaller cells in all organs than the larger adult flies (control/female individuals). Coupling between body size and cell size has been reported by previous studies [7,24,26,28,51,59], although this evidence was largely based on the measurements of single cell types (but see [8]). Systemic coordination of cell-size changes has been suggested by some interspecies [8,12,24,38,42] and intraspecies [20,39–41] comparisons, but until now, never demonstrated experimentally by manipulating the activity of cell-cycle regulatory pathways. Emerging evidence of the involvement of cell-cycle control in sex determination in *D. melanogaster* suggests that this control system includes autonomous regulation of cell size, which in females, but not in males, is additionally associated with a systemic signalling network via TOR/insulin pathways [43,60,61]. In light of our results, this sex-dependent regulatory system is tolerant to systemic changes in TOR pathway activity, maintaining sex differences in the cellular composition of organisms regardless of variation in TOR signalling activity. Indeed, Rideout et al. [60] showed that downregulation of TOR activity in larvae alone was not sufficient to alter sexual dimorphism of body size in adult *D. melanogaster*, whereas this dimorphism was blurred when the TOR and insulin pathways were simultaneously inhibited. Importantly, previous studies of sex-determination regulatory pathways in *Drosophila* have considered only single cell types; our results further suggest that while each cell in an organism maintains its sex identity by autonomously regulating its size, collectively, these regulations lead to highly orchestrated sexual differences in the cellular structure of tissues and organs. Overall, our evidence clearly shows that systemic regulation of cell size can be achieved via modulation of TOR activity. As part of the insulin/TOR signalling network, TOR is deactivated under natural conditions by a shortage of incoming nutrients and oxygen, which promotes autophagy and slows ageing [62,63]. This suggests that TOR plays a central role as a switch of resource allocation ‘sinks’, ultimately leading to the life-history strategy of an organism. In fact, *D. melanogaster* has evolved latitudinal clines in coupled changes in cell size, body size and genes affecting TOR activity [64–66], supporting the view that the activity of TOR and its phenotypic effects (cell size and body size) are not selectively neutral [13].

Taken together, our results and published data suggest that systemic orchestration of cell size and its contribution to the emergence of body-size variation occur commonly in nature, suggesting that these understudied phenomena are manifestations of adaptive responses to selection. Indeed, evidence suggests that *D. melanogaster* with rapamycin-induced reductions in the size of wing epidermal cells outperform flies with larger cells during flight in oxygen-poor conditions, suggesting a causal link between the cellularity of the body and organismal performance [29]. In support of this, we showed that changes in cell size in one cell type were associated with changes in cell size in other cell types, including flight muscles. However, it remains unclear how the collective effect of cell size in all tissues shapes organismal performance such that the synchronization of cell-size changes in different tissues confers evolutionary benefits. This is not a trivial question, especially as cell properties such as cell number, cell size, cell shape and organelle content should correspond closely to tissue-specific functions. The theory of optimal cell size (TOCS) [21,25,31,33,38,41,67,68] predicts that the cellular composition of an organism is optimized to selection pressures through a compromise between the cost of plasma-membrane maintenance and the cell capacity to perform physiological functions. The relatively large area of the plasma membrane of small cells should increase the rates of oxygen and nutrient fluxes but incurs costs imposed by ionic gradients and the need to maintain adequate membrane structure. In the TOCS framework, orchestration of the cellular composition of tissue throughout the body would maximize the benefits of having large or small cells, namely, by providing more efficient systemic energy savings or more efficient systemic transport of oxygen and nutrients, respectively. Certainly, maximizing the performance of highly specialized physiological functions, e.g., catabolic vs. anabolic processes, can require specific surface-to-volume ratios or organelle content, which could explain the reported irregularities in cell-size changes in different tissues [41].

In summary, we showed that organisms use developmental mechanisms to coordinate cell size in different organs and tissues and that this systemic cellular orchestration takes part in shaping the life-history strategy. For the first time, our study demonstrates the role of the TOR pathway in this cell-size coordination, which enables synchronization of the cellular composition of different organs in the body. Importantly, our results also suggest that the developmental sex-determination pathways involve tight coordination of cell size in different tissues of each sex, despite their cell-autonomous nature, as revealed by recent studies. This phenomenon deserves further investigation, especially because sexes often show different physiologies and life histories, including differences in longevity and susceptibility to different health issues [61]. We postulate that the activity of TOR/insulin pathways, with their systemic cellular effects, should be considered more frequently as part of various ecological and evolutionary patterns, such as the temperature-size rule (TSR) in ectotherms, Bergmann’s rule, Foster’s rule and Cope’s rule [9,69–73]. The incorporation of this activity can provide better understanding of the origins of fundamental biological phenomena, including sexual dimorphism, phylogenetic and geographical trends in life histories, and the developmental responses of ectotherms to climate change.

## Supporting information

Supplemental methods: histology & cell size measurements

## Supplementary material

The datasets, R code and materials supporting this article have been uploaded as part of the supplementary material.

## Authors’ contributions

E.S.: conceptualization, formal analysis, investigation, methodology, software, validation, visualization, writing — original draft, and writing — review & editing. A. M. L.: data curation, investigation, methodology, resources, supervision, validation, visualization, and writing – review & editing. V.P.: investigation, methodology, and writing – review & editing. A.S.: investigation, methodology, and writing – review & editing. M.C.: conceptualization, data curation, formal analysis, funding acquisition, methodology, project administration, resources, supervision, validation, visualization, writing – original draft, and writing – review & editing.

All authors gave final approval for publication and agreed to be held accountable for the work performed herein.

## References

1. McClain CR et al. 2015 Sizing ocean giants: Patterns of intraspecific size variation in marine megafauna. PeerJ, 3, e715. (doi:10.7717/peerj.715)

2. Watson W, Walker HJ. 2004 The world’s smallest vertebrate, *Schindleria brevipinguis*,a new paedomorphic species in the family Schindleriidae (Perciformes: Gobioidei). Rec. Aust. Museum 56, 139–142. (doi:10.3853/j.0067-1975.56.2004.1429)

3. Angilletta, Jr. MJ, Niewiarowski PH, Dunham AE, Leaché AD, Porter WP. 2004 Bergmann’s Clines in Ectotherms: Illustrating a Life-History Perspective with Sceloporine Lizards. Am. Nat. 164, E168–E183. (doi:10.1086/425222)

4. Zwaan BJ, Azevedo RBR, James AC, Van ‘T Land J, Partridge L. 2000 Cellular of wing size variation in *Drosophila melanogaster*: A comparison of latitudinal clines on two continents. Heredity 84, 338–347. (doi:10.1046/j.1365-2540.2000.00677.x)

5. Heinze J, Foitzik S, Fischer B, Wanke T, Kipyatkov VE. 2003 The significance of latitudinal variation in body size in a holarctic ant, *Leptothorax acervorum*. Ecography 26, 349–355. (doi:10.1034/j.1600-0587.2003.03478.x)

6. Partridge L, French V. 1996 Thermal evolution of ectotherm body size: why get big in the cold? In Animals and Temperature: Phenotypic and Evolutionary Adaptation (eds IA Johnston, AFE Bennett), pp. 265–292. Cambridge, UK: Cambridge University Press. (doi:10.1017/CBO9780511721854.012)

7. Partridge L, Langelan R, Fowler K, Zwaan B, French V. 1999 Correlated responses to selection on body size in *Drosophila melanogaster*. Genet. Res. 74, 43–54. (doi:10.1017/S0016672399003778)

8. Schramm BW, Labecka AM, Gudowska A, Antoł A, Sikorska A, Szabla N, Bauchinger U, Kozlowski J, Czarnoleski M. 2021 Concerted evolution of body mass, cell size and metabolic rate among carabid beetles. J. Insect Physiol. 132, 104272. (doi:10.1016/j.jinsphys.2021.104272)

9. Atkinson D. 1994 Temperature and Organism Size—A Biological Law for Ectotherms? Adv. Ecol. Res. 25, 1–58. (doi:10.1016/S0065-2504(08)60212-3)

10. Partridge L, Barrie B, Fowler K, French V. 1994 Evolution and Development of Body Size and Cell Size in *Drosophila melanogaster* in Response to Temperature. Evolution 48, 1269–1276. (doi:10.2307/2410384)

11. Klok CJ, Harrison JF. 2013 The temperature size rule in arthropods: Independent of macro-environmental variables but size dependent. Integr. Comp. Biol. 53, 557–570. (doi:10.1093/icb/ict075)

12. Heinrich EC, Farzin M, Klok CJ, Harrison JF. 2011 The effect of developmental stage on the sensitivity of cell and body size to hypoxia in *Drosophila melanogaster*. J. Exp. Biol. 214, 1419–1427. (doi:10.1242/jeb.051904)

13. Kozlowski J, Konarzewski M, Czarnoleski M. 2020 Coevolution of body size and metabolic rate in vertebrates: a life-history perspective. Biol. Rev. 95, 1393–1417. (doi:10.1111/brv.12615)

14. Harrison JF, Kaiser A, VandenBrooks JM. 2010 Atmospheric oxygen level and the evolution of insect body size. Proc. R. Soc. B Biol. Sci. 277, 1937–1946. (doi:10.1098/rspb.2010.0001)

15. Angilletta MJ, Sears MW. 2004 Evolution of thermal reaction norms for growth rate and body size in ectotherms: An introduction to the symposium. Integr. Comp. Biol. 44, 401–402. (doi:10.1093/icb/44.6.401)

16. Atkinson D, Sibly RM. 1997 Why are organisms usually bigger in colder environments? Making sense of a life history puzzle. Trends Ecol. Evol. 12, 235–239. (doi:10.1016/S0169-5347(97)01058-6)

17. Chown SL, Gaston KJ. 2010 Body size variation in insects: A macroecological perspective. Biol. Rev. 85, 139–169. (doi:10.1111/j.1469-185X.2009.00097.x)

18. Schramm BW, Gudowska A, Antoł A, Labecka AM, Bauchinger U, Kozlowski J, Czarnoleski M. 2018 Effects of fat and exoskeletal mass on the mass scaling of metabolism in Carabidae beetles. J. Insect Physiol. 106, 232–238. (doi:10.1016/j.jinsphys.2017.10.002)

19. Arendt J. 2007 Ecological correlates of body size in relation to cell size and cell number: Patterns in flies, fish, fruits and foliage. Biol. Rev. 82, 241–256. (doi:10.1111/j.1469-185X.2007.00013.x)

20. Azevedo RBR, French V, Partridge L. 2002 Temperature modulates epidermal cell size in *Drosophila melanogaster*. J. Insect Physiol. 48, 231–237. (doi:10.1016/S0022-1910(01)00168-8)

21. Czarnoleski M, Dragosz-Kluska D, Angilletta, Jr. MJ. 2015 Flies developed smaller cells when temperature fluctuated more frequently. J. Therm. Biol. 54, 106–110. (doi:10.1016/j.jtherbio.2014.09.010)

22. Antoł A, Labecka AM, Horváthová T, Zieliński B, Szabla N, Vasko Y, Pecio A, Kozlowski J, Czarnoleski M. 2020 Thermal and oxygen conditions during development cause common rough woodlice (*Porcellio scaber*) to alter the size of their gas-exchange organs. J. Therm. Biol. 90, 102600. (doi:10.1016/j.jtherbio.2020.102600)

23. Vijendravarma RK, Narasimha S, Kawecki TJ. 2011 Plastic and evolutionary responses of cell size and number to larval malnutrition in *Drosophila melanogaster*. J. Evol. Biol. 24, 897–903. (doi:10.1111/j.1420-9101.2010.02225.x)

24. Kozlowski J, Czarnoleski M, François-Krassowska A, Maciak S, Pis T. 2010 Cell size is positively correlated between different tissues in passerine birds and amphibians, but not necessarily in mammals. Biol. Lett. 6, 792–796. (doi:10.1098/rsbl.2010.0288)

25. Czarnoleski M, Labecka AM, Starostová Z, Sikorska A, Bonda-Ostaszewska E, Woch K, Kubička L, Kratochvíl L, Kozlowski J. 2017 Not all cells are equal: Effects of temperature and sex on the size of different cell types in the Madagascar ground gecko *Paroedura picta*. Biol. Open 6, 1149–1154. (doi:10.1242/bio.025817)

26. Czarnoleski M, Cooper BS, Kierat J, Angilletta, Jr. MJ. 2013 Flies developed small bodies and small cells in warm and in thermally fluctuating environments. J. Exp. Biol. 216, 2896–2901. (doi:10.1242/jeb.083535)

27. Adrian GJ, Czarnoleski M, Angilletta MJ. 2016 Flies evolved small bodies and cells at high or fluctuating temperatures. Ecol. Evol. 6, 7991–7996. (doi:10.1002/ece3.2534)

28. Starostová Z, Kratochvíl L, Frynta D. 2005 Dwarf and giant geckos from the cellular perspective: The bigger the animal, the bigger its erythrocytes? Funct. Ecol. 19, 744–749. (doi:10.1111/j.1365-2435.2005.01020.x)

29. Szlachcic E, Czarnoleski M. 2021 Thermal and oxygen flight sensitivity in ageing *Drosophila melanogaster* flies: Links to rapamycin-induced cell size changes. Biology 10, 861. (doi:10.3390/biology10090861)

30. Verspagen N, Leiva FP, Janssen IM, Verberk WCEP. 2020 Effects of developmental plasticity on heat tolerance may be mediated by changes in cell size in *Drosophila melanogaster*. Insect Sci. 27, 1244–1256. (doi:10.1111/1744-7917.12742)

31. Walczyńska A, Labecka AM, Sobczyk M, Czarnoleski M, Kozlowski J. 2015 The Temperature-Size Rule in *Lecane inermis* (Rotifera) is adaptive and driven by nuclei size adjustment to temperature and oxygen combinations. J. Therm. Biol. 54, 78–85. (doi:10.1016/j.jtherbio.2014.11.002)

32. Verberk WCEP, Sandker JF, van de Pol ILE, Urbina MA, Wilson RW, McKenzie DJ, Leiva FP. 2022 Body mass and cell size shape the tolerance of fishes to low oxygen in a temperature-dependent manner. Glob. Chang. Biol. 28, 5695–5707. (doi:10.1111/gcb.16319)

33. Liu S, Tan C, Tyers M, Zetterberg A, Kafri R. 2022 What programs the size of animal cells? Front. Cell Dev. Biol. 10, 949382. (doi:10.3389/fcell.2022.949382)

34. Schramm BW, Gudowska A, Kapustka F, Labecka AM, Czarnoleski M, Kozlowski J. 2015 Automated measurement of ommatidia in the compound eyes of beetles. Biotechniques 59, 99–101. (doi:10.2144/000114316)

35. Maciak S, Janko K, Kotusz J, Choleva L, Boroń A, Juchno D, Kujawa R, Kozlowski J, Konarzewski M. 2011 Standard Metabolic Rate (SMR) is inversely related to erythrocyte and genome size in allopolyploid fish of the *Cobitis taenia* hybrid complex. Funct. Ecol. 25, 1072–1078. (doi:10.1111/j.1365-2435.2011.01870.x)

36. Chown SL, Marais E, Terblanche JS, Klok CJ, Lighton JRB, Blackburn TM. 2007 Scaling of insect metabolic rate is inconsistent with the nutrient supply network model. Funct. Ecol. 21, 282–290. (doi:10.1111/j.1365-2435.2007.01245.x)

37. Starostová Z, Konarzewski M, Kozlowski J, Kratochvíl L. 2013 Ontogeny of Metabolic Rate and Red Blood Cell Size in Eyelid Geckos: Species Follow Different Paths. PLoS One 8, 21–23. (doi:10.1371/journal.pone.0064715)

38. Czarnoleski M, Labecka AM, Dragosz-Kluska D, Pis T, Pawlik K, Kapustka F, Kilarski WM, Kozlowski J. 2018 Concerted evolution of body mass and cell size: Similar patterns among species of birds (Galliformes) and mammals (Rodentia). Biol. Open 7, bio029603. (doi:10.1242/bio.029603)

39. Antoł A, Labecka AM, Horváthová T, Sikorska A, Szabla N, Bauchinger U, Kozlowski J, Czarnoleski M. 2020 Effects of thermal and oxygen conditions during development on cell size in the common rough woodlice *Porcellio scaber*. Ecol. Evol. 10, 9552–9566. (doi:10.1002/ece3.6683)

40. Stevenson RD, Hill MF, Bryant PJ. 1995 Organ and cell allometry in Hawaiian *Drosophila*: How to make a big fly. Proc. R. Soc. B Biol. Sci. 259, 105–110. (doi:10.1098/rspb.1995.0016)

41. Maciak S, Bonda-Ostaszewska E, Czarnoleski M, Konarzewski M, Kozlowski J. 2014 Mice divergently selected for high and low basal metabolic rates evolved different cell size and organ mass. J. Evol. Biol. 27, 478–487. (doi:10.1111/jeb.12306)

42. Brodribb TJ, Jordan GJ, Carpenter RJ. 2013 Unified changes in cell size permit coordinated leaf evolution. New Phytol. 199, 559–570. (doi:10.1111/nph.12300)

43. Deng H, Jasper H. 2016 Sexual Dimorphism: How Female Cells Win the Race. Curr. Biol. 26, R212–R215. (doi:10.1016/j.cub.2016.01.062)

44. Roustan V, Jain A, Teige M, Ebersberger I, Weckwerth W. 2016 An evolutionary perspective of AMPK–TOR signaling in the three domains of life. J. Exp. Bot. 67, 3897–3907. (doi:10.1093/jxb/erw211)

45. van Dam TJP, Zwartkruis FJT, Bos JL, Snel B. 2011 Evolution of the TOR Pathway. J. Mol. Evol. 73, 209–220. (doi:10.1007/s00239-011-9469-9)

46. Bjedov I, Toivonen JM, Kerr F, Slack C, Jacobson J, Foley A, Partridge L. 2010 Mechanisms of Life Span Extension by Rapamycin in the Fruit Fly *Drosophila melanogaster*. Cell Metab. 11, 35–46. (doi:10.1016/j.cmet.2009.11.010)

47. Rohde PD, Bøcker A, Jensen CAB, Bergstrøm AL, Madsen MIJ, Christensen SL, Villadsen SB, Kristensen TN. 2021 Genotype and Trait Specific Responses to Rapamycin Intake in *Drosophila melanogaster*. Insects 12, 474. (doi:doi.org/10.3390/insects12050474)

48. Li Z, Zhang Z, Ren Y, Wang Y, Fang J, Yue H, Ma S, Guan F. 2021 Aging and age-related diseases: from mechanisms to therapeutic strategies. Biogerontology 22, 165–187. (doi:10.1007/s10522-021-09910-5)

49. Wu MYW, Cully M, Andersen D, Leevers SJ. 2007 Insulin delays the progression of *Drosophila* cells through G2/M by activating the dTOR/dRaptor complex. EMBO J. 26, 371–379. (doi:10.1038/sj.emboj.7601487)

50. Potter S, Sifers J, Yocom E, Blümich SLE, Potter R, Nadolski J, Harrison DA, Cooper RL. 2019 Effects of inhibiting mTOR with rapamycin on behavior, development, neuromuscular physiology and cardiac function in larval *Drosophila*. Biol. Open 8, bio046508. (doi:10.1242/bio.046508)

51. Czarnoleski M, Labecka AM, Kozlowski J. 2016 Thermal plasticity of body size and cell size in snails from two subspecies of *Cornu aspersum*. J. Molluscan Stud. 82, 235–243. (doi:10.1093/mollus/eyv059)

52. R Core Team. 2020 R: A language and environment for statistical computing. R Foundation for Statistical Computing, Vienna, Austria. (URL https://www.R-project.org/)

53. Bates D, Mächler M, Bolker BM, Walker SC. 2015 Fitting linear mixed-effects models using lme4. J. Stat. Softw. 67, 1–48. (doi:10.18637/jss.v067.i01)

54. Kuznetsova A, Brockhoff PB, Christensen RHB. 2017 lmerTest Package: Tests in Linear Mixed Effects Models. J. Stat. Softw. 82, 1–26. (doi:10.18637/jss.v082.i13)

55. Fox J, Weisberg S. 2019 An R Companion to Applied Regression. Thousand Oaks, USA: SAGE Publications, Inc (https://socialsciences.mcmaster.ca/jfox/Books/Companion/)

56. Wickham H. 2016 ggplot2 Elegant Graphics for Data Analysis. New York, NY, USA: Springer.

57. Lenth R V. 2020 emmeans: Estimated Marginal Means, aka Least-Squares Means. R package version 1.5.3.

58. Harrington B, Gould T, Hurst N, MenTaLgu Y. 2004-2005 Inkscape. (https://www.inkscape.org)

59. Davison J. 1956 An analysis of cell growth and metabolism in the crayfish (*Procambarus allenĩ*). Biol. Bull. 110, 264–273. (doi:10.2307/1538832)

60. Rideout EJ, Narsaiya MS, Grewal SS. 2016 The Sex Determination Gene transformer Regulates Male-Female Differences in *Drosophila* Body Size. PLOS Genet. 11, e1005683. (doi:10.1371/journal.pgen.1005683)

61. Hudry B, Khadayate S, Miguel-Aliaga I. 2016 The sexual identity of adult intestinal stem cells controls organ size and plasticity. Nature 530, 344–348. (doi:10.1038/nature16953)

62. Grewal SS. 2009 Insulin/TOR signaling in growth and homeostasis: A view from the fly world. Int. J. Biochem. Cell Biol. 41, 1006–1010. (doi:10.1016/j.biocel.2008.10.010)

63. Miki T, Shinohara T, Chafino S, Noji S, Tomioka K. 2020 Photoperiod and temperature separately regulate nymphal development through JH and insulin/TOR signaling pathways in an insect. Proc. Natl. Acad. Sci. U. S. A. 117, 5525–5531. (doi:10.1073/pnas.1922747117)

64. De Jong G, Bochdanovits Z. 2003 Latitudinal clines in *Drosophila melanogaster:* Body size, allozyme frequencies, inversion frequencies, and the insulin-signalling pathway. J. Genet. 82, 207–223. (doi:10.1007/BF02715819)

65. Paaby AB, Blacket MJ, Hoffmann AA, Schmidt PS. 2010 Identification of a candidate adaptive polymorphism for *Drosophila* life history by parallel independent clines on two continents. Mol. Ecol. 19, 760–774. (doi:10.1111/j.1365-294X.2009.04508.x)

66. Fabian DK, Kapun M, Nolte V, Kofler R, Schmidt PS, Schlötterer C, Flatt T. 2012 Genome-wide patterns of latitudinal differentiation among populations of *Drosophila melanogaster* from North America. Mol. Ecol. 21, 4748–4769. (doi:10.1111/j.1365-294X.2012.05731.x)

67. Hermaniuk A, Van De Pol ILE, Verberk WCEP. 2021 Are acute and acclimated thermal effects on metabolic rate modulated by cell size? A comparison between diploid and triploid zebrafish larvae. J. Exp. Biol. 224, jeb227124. (doi:10.1242/jeb.227124)

68. Szarski H. 1983 Cell size and the concept of wasteful and frugal evolutionary strategies. J. Theor. Biol. 105, 201–209. (doi:10.1016/S0022-5193(83)80002-2)

69. Van Voorhies WA. 1996 Bergmann size clines: A simple explanation for their occurrence in ectotherms. Evolution 50, 1259–1264. (doi:10.1111/j.1558-5646.1996.tb02366.x)

70. Van Valen L. 1973 Body Size and Numbers of Plants and Animals. Evolution 27, 27–35. (doi:10.2307/2407116)

71. Lokatis S, Jeschke JM. 2018 The island rule: An assessment of biases and research trends. J. Biogeogr. 45, 289–303. (doi:10.1111/jbi.13160)

72. Stanley SM. 1973 An Explanation for Cope’s Rule. Evolution 27, 1–26. (doi:10.2307/2407115)

73. Rensch B. 1948 Histological Changes Correlated with Evolutionary Changes of Body Size. Evolution 2, 218–230. (doi:10.2307/2405381)

