## Supplemental methods: histology & cell size measurements for "Systemic orchestration of cell size throughout the body: Influence of sex and rapamycin exposure in *Drosophila melanogaster*"

**Supplementary material**

After dissection, wings were preserved at -20°C in Eppendorf tubes. For measurements, wings were mounted on microscopy slides using ST Ultra and CV Ultra (Leica Biosystems, Nussloch, Germany) and digitalized under 20× objective magnification. According to Dobzhansky [1], each trichome represents one wing epidermal cell. Thus, we calculated the mean wing epidermal cell size (μm$^2$) from the ratio of defined wing area and the number of corresponding trichomes. Based on previous methods [2,3], we counted trichomes on the dorsal wing blade in a 0.031-mm$^2$ circle placed between the cubital and distal veins with the help of two macros embedded in ImageJ software (National Institutes of Health (NIH), Bethesda, MD, USA).

Heads were preserved with 1 ml of 20% methanol (POCH, Gliwice, Poland). Ommatidia, which form compound eyes in *Drosophila*, are composed of a fixed number of cells [4]. Thus, based on previous studies [5-7], we used their size as a proxy of cell size. To conduct the measurements, both eyes were covered with a transparent 5SecondFix UV glue and dried with a UV lamp (Ontel Products, Fairfield, USA). Such prepared replicas were removed from the eyes and flattened on a slide by incising in three places. Next, a drop of glycerine (POCH) was added, and the slide was covered with a coverslip. On the top of each slide, fishing weights (100 g) were placed for 24 h to allow flattening of replicas, and then the coverslip edges were covered with colourless nail polish (Vipera, Kraków, Poland). Prepared slides were digitalized under 10× objective magnification. We estimated the mean ommatidium size (μm$^2$) using automatized methods [5] integrated into ImageJ (NIH).

Legs were preserved with 1 ml of 70% ethanol (Linegal Chemicals, Warszawa, Poland). For epidermal cell measurements, legs were placed individually on microscopic glass slides to obtain photos of the 8th row of the basitarsus under 20× objective magnification. Using ZEN software (Zeiss), we measured the distance (μm) between the first and last bristles of the 8th row (based on Hannah-Alava nomenclature [8]) and counted the total number of bristles. According to Held [9], the bristle number in a row and cell number along the basitarsus are coordinated as are the bristle interval and cell diameter. Therefore, to calculate the proximal cell size (μm), the measured distance was divided by the bristle number in the row.

During dissection, the Malpighian tubules were taken with two entomological pins (Paradox, Kraków, Poland) from the severed abdomen on the microscopic glass with a drop of PBSTX (mixture of phosphate-buffered saline (PBS; Sigma Aldrich, Saint Louis, USA) and Triton X-100 (POCH)). After preparation, slides were covered with a coverslip, and Malpighian tubules were digitalized under 20× magnification with a phase contrast objective. We estimated the proxy of cell size forming Malpighian tubules ($\mu m^2$) from the density of cell nuclei/nucleoli in a known area of the tubule. We outlined the area of Malpighian tubules using ImageJ software with a LiveWire Plugin (NIH) and counted the number of cell nuclei/nucleoli in this area. We carried out measurements in five to ten photos for each individual.

Dissected thoraxes were used to measure dorsal longitudinal muscle cells, a subset of indirect flight muscle cells. According to the modified methodology of Barbosa et al. [10], to fix this tissue, thoraxes were placed in Duboscq–Brasil's solution for 2 h. Within this time, Eppendorf tubes with thoraxes were placed in a beaker with water standing on a hot plate (50 °C) for 10 minutes. Next, thoraxes were washed in PBSTX, dehydrated in 70%, 80%, 90% and 96% ethanol (Linegal Chemicals), butanol (Chempur, Piekary Śląskie, Poland) and isopropanol (Leica), cleared in pure Clearene (Leica), then in a mixture of Clearene (Leica) and Paraplast Plus (Leica) and finally embedded in Paraplast Plus (Leica). Thoraxes were cut with a rotary microtome (Hyrax M55, Zeiss) into 6-μm thick cross-sections. The slides were deparaffinized in Clearene (Leica), rehydrated in isopropanol (Leica) and 96%, 90%, 80%, 70% and 50% ethanol (Linegal) and distilled water, and stained with Ehrlich haematoxylin (Carl Roth) and then with Gömöri trichrome (BioOptica, Milano, Italy). Then, the slides were washed in 0.2% acetic acid (POCH), dehydrated in 96% ethanol (Linegal), cleared in a mixture of phenol (Chempur) and xylene (POCH) and then in pure xylene (POCH), and embedded in CV Mount (Leica). We analysed muscle tissue under 20× objective magnification and photographed one bundle from each of the six pairs of muscle bundles. Using ImageJ (NIH), we outlined the area of individual muscle cells (fibres) ($\mu m^2$). We measured 60 cells for each fly.

### References


1. Dobzhansky T. 1929 The influence of the quantity and quality of chromosomal material on the size of the cells in *Drosophila melanogaster*. *Wilhelm Roux Arch. Entwickl. Mech. Org.* **115**, 363–379. (doi:10.1007/BF02078996)

2. Czarnoleski M, Dragosz-Kluska D, Angilletta, Jr. MJ. 2015 Flies developed smaller cells when temperature fluctuated more frequently. *J. Therm. Biol.* **54**, 106–110. (doi:10.1016/j.jtherbio.2014.09.010)

3. Szlachcic E, Czarnoleski M. 2021 Thermal and oxygen flight sensitivity in ageing *Drosophila melanogaster* flies: Links to rapamycin-induced cell size changes. *Biology* **10**, 861. (doi:10.3390/biology10090861)

4. Cagan R. 2009 Principles of *Drosophila* eye differentiation. *Curr. Top. Dev. Biol.* **89**, 115–135. (doi:10.1016/S0070-2153(09)89005-4)

5. Schramm BW, Gudowska A, Kapustka F, Labecka AM, Czarnoleski M, Kozlowski J. 2015 Automated measurement of ommatidia in the compound eyes of beetles. *Biotechniques* **59**, 99–101. (doi:10.2144/000114316)

6. Kramer JM, Davidge JT, Lockyer JM, Staveley BE. 2003 Expression of *Drosophila* FOXO regulates growth and can phenocopy starvation. *BMC Dev. Biol.* **3**, 1471–213X/3/5. (doi:10.1186/1471-213X-3-5)

7. Montagne J, Stewart MJ, Stocker H, Hafen E, Kozma SC, Thomas G. 1999 *Drosophila* S6 kinase: A regulator of cell size. *Science* **285**, 2126–2129. (doi:10.1126/science.285.5436.2126)

8. Hannah-Alava A. 1958 Morphology and chaeotaxy of the legs of *Drosophila melanogaster*. *J Morphol.* **103**, 281–310. (doi:org/10.1002/jmor.1051030205)

9. Held LI. 1979 Pattern as a function of cell number and cell size on the second-leg basitarsus of *Drosophila*. *Wilhelm Roux Arch. Dev. Biol.* **187**, 105–127. (doi:10.1007/BF00848266)

10. Barbosa P, Berry D, Kary CK. 2015 *Insect Histology – Practical Laboratory Techniques*. Hoboken, NJ, USA: Willey Blackwell.
